# Acute exposure of fruit bats to low concentrations of heavy metals affects oxidative stress markers

**DOI:** 10.1101/2023.06.15.545099

**Authors:** Ana Luiza Fonseca Destro, Thaís Silva Alves, Fernanda Ribeiro Dias, Reggiani Vilela Gonçalves, Jerusa Maria de Oliveira, Leandro Licursi de Oliveira, Mariella Bontempo Freitas

**Affiliations:** Department of Animal Biology, Federal University of Viçosa, Viçosa, MG, Brazil; Department of Veterinary Medicine, Federal Rural University of Pernambuco, Recife, PE, Brazil; Institute of Chemistry and Biotechnology, Federal University of Alagoas, Maceió, AL, Brazil; Department of General Biology, Federal University of Viçosa, Viçosa, MG, Brazil

**Keywords:** Chiroptera, *Artibeus lituratus*, Ecotoxicology, Integrated Biomarker Response

## Abstract

Cadmium (Cd), chromium (Cr), lead (Pb), and nickel (Ni) are heavy metals and common environmental pollutants. We aimed to investigate heavy metals’ effects on fruit-bats’ organs. Adult males (*Artibeus lituratus*) were captured and exposed to heavy metals (1.5 mg/kg). The Integrated Biomarker Response helped us understand the interrelationship in a multi-biomarker global approach to oxidative stress. The liver was more sensitive to Ni and Pb than Cd and Cr. In the kidney, Pb did not cause hazardous effects, unlike the other metals. In testes, Ni doubled damage levels compared to the other metals. Ni did not cause serious effects in muscles, which was more sensitive to Pb and Cd than to Cr. The brain was more susceptible to Pb and Ni than Cr and Cd. We observed that acute doses, even in low concentrations, are deleterious to fruit-bats. We propose the following order of metal toxicity: Ni> Pb> Cd> Cr.

**SUMMARY STATEMENT:** The investigation of heavy metals toxicity in fruit bats reveals differential sensitivities of organ and highlights the harmful effects of acute exhibitions even at low concentrations.

## INTRODUCTION

In ecotoxicological studies, heavy metals (HM) are often referred to as metals and metalloids with potential toxicity, usually associated with environmental pollution (Ali et al., 2019). Anthropogenic activities such as mining and industrialization have been a major concern as they might be involved in increased HM levels in the environment (Ali et al., 2013). As metals are not degraded in the environment, they may accumulate in living organisms and within food chains, processes known as bioaccumulation and biomagnification, respectively (Ali et al., 2013).

Cadmium (Cd) is a non-essential HM (Mustafa and Mustafa, 2000) and a common persistent environmental pollutant. It can accumulate in organisms through food chains (Liu et al., 2020). In the male mammalian reproductive system, Cd exposure shows one of the main disruptive potentials among HM (Tariba Lovaković, 2020), causing hormone and sperm alterations (da Silva et al., 2021; Güner et al., 2020; Tariba Lovaković, 2020). Low concentrations and prolonged exposure to Cd also damaged different organs, such as the liver and kidneys (Genchi et al., 2020). Chromium (Cr) is naturally found in soil and water. It is an essential animal nutrient (Shrivastava et al., 2002), and it has been heavily used in metallurgical, chemical, and refractory industries (Chandra et al., 2007). Excessive Cr exposure affects the mammalian reproductive system, inducing oxidative stress due to its strong oxidant capacity (Chandra et al., 2007). Prolonged exposures also cause genotoxic effects and disturbances to the immune system (Shrivastava et al., 2002). Lead (Pb) is a non-essential element (Mustafa and Mustafa, 2000) widely found in industrial products (batteries, paints, gasoline, pesticides, medicines, cosmetics, etc.). It is considered the most important toxic HM in the environment due to its abundant global distribution (Wani et al., 2015). Pb induces damage to renal, reproductive, and nervous systems in mammals through excessive reactive oxygen and nitrogen species (ROS/NOS) production and antioxidant enzyme inhibition (Fan et al., 2020; Wani et al., 2015). Nickel (Ni) has physical and chemical properties that favour its use in stainless steel manufacturing, structural steel alloys, electroplating, and batteries (Kong et al., 2021). It is an essential nutrient (Genchi et al., 2020), however, high Ni levels have been proven to disrupt the Hypothalamus-Pituitary-Gonad (HPG) axis and generate excessive ROS/NOS production in testes (Forgacs et al., 2012).

Besides, mining activities are also involved with eventual dam rupture, as recently happened twice in Brazil (in the Atlantic Forest area), and might as well increase the associated metal concentration in the environment. These associated metals are often not bioavailable, although they can be released due to anoxic conditions associated with plant and animal activity (Queiroz et al., 2018). In Brazilian tropical basins, these metals were found in sediments, in concentrations (mg/kg) varying from below the detection limit to 6.6 for Cd, 2.2 to 119.5 for Cr, and below the detection limit to 92 for Pb (Quadra et al., 2019). This environmental contamination may impact the local fauna in several ways and the extent depends on other factors. Probably, the animals’ trophic level influences the HM impact extent, as other factors like the chemical form of the metal in the environment, availability, concentration, exposure time, and interaction with other metals (Hejna et al., 2018). Although the effects of HM have been described in several animal models, its impact on wild species, constantly exposed to them orally and through skin contact, is less understood. Bats are the only true flying mammals, with adaptations that include a low reproduction rate (Barclay et al., 2004), high metabolic rate, high longevity, and a reduced inflammatory response (may be associated with their ability to deal with viral infection with lower impact (Moreno Santillán et al., 2021). Some of these specific traits could indicate that environmental pollutants affect bats differently than other mammals (Secord et al., 2015). Despite this, the risk assessment of contaminants in bats is developing and requires a specific approach for these animals (Hernández-Jerez et al., 2019). Bats also play a vital role in ecosystem services through pollination (bats pollinate 549 plant species), pest control (752 insect species are consumed by bats, including crop pests and disease vectors), and seed dispersal (Ramírez-Fráncel et al., 2022). The great fruit-eating bat (*Artibeus lituratus*) is an abundant species in the Neotropical region. It contributes to reforestation, especially in heavily fragmented areas of the Atlantic Rain Forest (Laurindo and Vizentin-Bugoni, 2020; Oliveira et al., 2021). In these regions, though, fruit bats face several challenges to survive, including HM pollution (Zukal et al., 2015).

Despite a few limitations, bats are considered potential ecological bioindicators, mainly due to their ability to inhabit wide geographic ranges, to be at the top of the food chain, to show ecological and evolutionary interactions with other ecosystem components, such as bat-insect and bat-plant coevolution, and for the vital ecosystem services provided (Russo et al., 2021). A few *in loco* toxicological studies associate environmental exposure to HMs, detected in guano or blood, with damage to the liver, DNA, and cholinergic functions (Zukal et al., 2015). The scientific literature shows that bats can accumulate HMs found in the environment. Concentrations (mg/kg) such as 5.8 – 7.32 of Pb, 5.7 – 10.9 of Cr, 3.6 – 4.05 of Cd, and 4.3 – 8.6 of Ni have already been found in insectivorous bats living at coal mining areas (José Zocche et al., 2010). However, there is a lack of information on the specific, isolated effect of the main environmentally available HMs in bats exposed to each one individually. In a region where mining operations are close to forests, investigations on how HM pollution affects essential aspects of local wildlife are needed to understand the extension of these impacts.

Here we aimed to evaluate the toxicological effects of short-term exposure to lower concentrations than found in the environment of four important HMs (Cd, Cr, Pb, and Ni) on oxidative parameters in physiologically relevant tissues in the great fruit-eating bat.

## MATERIAL AND METHODS

### Chemical

The metals CdCl_2_ (cadmium chloride 99.9%), CrO_3_ (chromium VI trioxide ≥99%), Pb (CH_3_COO)^2^·3H_2_O (neutral lead acetate PA trihydrate), and Cl_2_Ni.6H_2_O (nickel (II) chloride hexahydrate 99.9%) were obtained from Sigma-Aldrich (St. Louis, Missouri, US) and Merck (Darmstadt, Germany).

### Animals

Adult male great fruit-eating bats (*Artibeus lituratus*, n=31, BW=73.62±5.78 g) were captured using mist nets in a forest area from the Federal University of Viçosa (UFV) (20° 45’ S and 42° 52’ W), Viçosa, Minas Gerais, Brazil. According to Díaz (Díaz et al., 2016), all animals were identified and brought to the University and kept in a half-wall screen-lined bat house located at the Museum of Zoology garden, under a few trees of the Atlantic Forest. Bats were assigned to individual enclosures (8 of 2 m^3^ each). They could fly freely inside the rooms and keep under natural temperature, light, and humidity cycles. All animals were submitted to a 4-day acclimation period before the exposure to HM started. During this time, the animals were offered tropical fruits (*Carica papaya, Musa* sp, *Psidium guajava*, and *Mangifera indica* L.) and water *ad libitum*. All procedures performed in this study were approved by the Brazilian Government (SISBIO 75064-1) and by the Animal Ethics Committee (CEUA-UFV 26/2020). The approval certificates are available upon request.

### Experimental design

Following the acclimation period, the animals were treated according to one of the five experimental groups: CTL) control: saline solution (NaCl 0.9%) (n=6) and experimental groups that received 1.5 mg/kg of each metal: Cd) cadmium chloride (CdCl_2)_ (n=6); Cr) chromium trioxide IV (CrO_3)_ (n=6); Pb) lead acetate (Pb (CH_3_COO)_2_·3H_2_O) (n=6) and Ni) nickel chloride (Cl_2_Ni.6H_2_O) (n=7). Exposure to saline or metals was performed through one intraperitoneal injection (i.p.) of 0.7 mL of solution at 8:00 am on day 1 of exposure). The i.p. route was chosen to ensure that the total volume would be entirely absorbed by the animal, avoiding consumption bias between metals (Al Shoyaib et al., 2020). Food (fruits) was offered each night at 6:00 pm (150-200 g for each bat). According to Oliveira (Oliveira et al., 2021), leftovers were weighed in the morning to ensure that all animals were fed to satisfaction. Water was available *ad libitum*. The metal concentrations we tested were chosen from previous similar experiments with adult male mice (Cupertino et al., 2017; Mouro et al., 2020). Although LC50 doses are already stipulated for murine models, this study is the first to use bats as models. The effects of each metal on bats are unclear such as his concentration-time effects on these animals. The same concentrations for all metals were chosen to make comparisons among them possible. After 96h of exposure, bats were euthanized through cervical dislocation followed by decapitation. Blood was collected from the trunk in tubes containing heparin for further biochemical markers determination. The liver, kidney, breast muscle, testes, and brain were rapidly removed under ice, divided into fragments, and weighed. Portions assigned to the redox status determination were frozen in liquid nitrogen until storage at -80°C.

### Biochemical biomarkers in plasma

Blood samples (∼3.5 mL) were centrifuged at 1500*g* for 15min. The plasma was separated in aliquots of 0.5 mL and stored at −80 °C until the determination of <u>aspartate</u> aminotransferase (AST), alanine aminotransferase (ALT), triglycerides, cholesterol, glucose, and albumin. All parameters were measured through enzymatic colourimetry using a commercial kit (Bioclin, Belo Horizonte, Brazil).

### Tissue preparation

Samples were homogenized in 0.2 mol/L phosphate buffer and 1 mmol/L ethylenediaminetetraacetic acid (EDTA) (1:1 and 1.5:1, respectively), at a pH 7.4, using a tissue homogenizer (OMNI) (Kennesaw, USA). The homogenates were centrifuged at 15,000 g for 10 min at 4 °C before the analysis.

### Detoxification biomarkers

The homogenate supernatant was used to measure superoxide dismutase (SOD), catalase (CAT), glutathione S-transferase (GST), and ferric-reducing ability of plasma (FRAP). All samples were randomly assigned to blind analyses without sample identification until the analysis of the results to avoid bias. All samples were run in duplicates using a spectrophotometer (UV-Mini 1240, Shimadzu, Japan) or a microplate reader (Thermo Scientific, Waltham, USA).

SOD activity was determined based on the reduction of the superoxide (O^−2^) and hydrogen peroxide, thereby decreasing the auto-oxidation of pyrogallol (Dieterich et al., 2000). The reaction mixture contained 99 μL of potassium phosphate buffer (5 mmol/L, pH 8.0) and 30 μL of the sample. It was started by adding 15 μL of pyrogallol (100 μmol/L). The final reaction was measured by absorbance at 570 nm. SOD activity was calculated as units per milligram of protein. One unit (U) of SOD was defined as the amount that inhibited the rate of pyrogallol autoxidation by 50%. Duplicates of standards and blank samples for SOD activity were prepared with and without pyrogallol, respectively.

CAT activity was determined by adapting the Hadwan and Abed method (Hadwan and Abed, 2016). Briefly, 5 μL samples were incubated with 100 μL hydrogen peroxide (20 mmol/L), 100 μL of sodium, and potassium phosphate pH buffer (50 mmol/L, pH 7.0). After 3 min, the reaction was stopped with 150 μL ammonium molybdate (32.4 mmol/L). A control test without hydrogen peroxide was used to exclude the interference of amino acids and proteins. The reading, at 374 nm, was performed in a spectrophotometer. To calculate CAT activity, a standard curve was built with serial dilutions of hydrogen peroxide. CAT activity was expressed in CAT KU/milligrams of protein.

GST activity was measured using the method of Habig (Habig et al., 1974). Briefly, 1 mmol/L of glutathione-conjugated 1-chloro-2,4-dinitrochlorobenzene (CDNB) was added to the buffer containing one mmol/L of GSH and to an aliquot (10 μL) of the homogenate supernatant. Upon the addition of DNCB, the alteration was monitored through absorbance at 340 nm for 60 s. The molar extinction coefficient used for DNCB was ε340=9.6 mmol/L × cm. GST activity was expressed in μmol/min/g.

The total antioxidant capacity was estimated according to the ferric reducing antioxidant power (FRAP), a method described by Benzie and Strain using TPTZ (2,4,6-Tris(2-pyridyl)-s-triazine) as a substrate (Benzie and Strain, 1996). The method reduces a ferric 2,4,6-tripyridyl-s-triazine complex (Fe^3+^-TPTZ) to the ferrous form (Fe^2+^-TPTZ). Samples (10 μL) were added as FRAP solution (190 μL) of 25mL of acetate buffer (300 mmol/L, pH3.6), 2.5mL of TPTZ reagent (10 mmol/L), and 2.5 FeCl_3_.6H_2_O solutions (20 mmol/L) and the increase in absorbance at 593 nm was measured. The reducing Fe^3+^-TPTZ reagent by antioxidants was determined using the standard curve of serial dilutions of FeSO_4_·7H_2_O starting with 2 mmol/L. The FRAP was expressed as μM.

### Damage biomarkers

The homogenate supernatant was used to measure nitric oxide (NO), malondialdehyde (MDA), and protein carbonyl (PC). All samples were randomly assigned to blind analyses without sample identification until the analysis of the results to avoid bias. All samples were run in duplicates using a spectrophotometer (UV-Mini 1240, Shimadzu, Japan) or a microplate reader (Thermo Scientific, Waltham, USA).

The standard Griess reaction quantified NO production. Briefly, 50 μL of supernatants described above were incubated with an equal volume of Griess reagent (1% sulfanilamide, 0.1% N-(1-Naphthyl) ethylenediamine and 2.5% phosphoric acid) at room temperature for 10 min (Tsikas, 2007). The absorbance was measured at 570 nm in a microplate reader. The conversion of absorbance into micromolar concentrations of NO was obtained from a sodium nitrite (0–100 μmol/L) standard curve and expressed as NO concentrations (μmol/L).

Malondialdehyde (MDA) is the major product of lipid peroxidation. MDA was measured according to Buege and Aust (Buege and Aust, 1978). Briefly, 0.2 mL of the tissue supernatant was homogenized in a solution (0.4 mL) of trichloroacetic acid (15%)/thiobarbituric acid (0.375%)/hydrochloric acid (0.6%). The total reaction mixture was kept in a boiling water bath for 40 min. After cooling on ice, butyl alcohol (0.6 mL) was added. The solution was vortexed for 2 minutes and centrifuged for 10 minutes at 9,000 g. The supernatant was used to measure the absorbance at 535 nm. The concentration of MDA was determined by using the standard curve of known concentrations of 1,1,3,3-tetramethoxypropane (TEMPO). The results were expressed as μmol/ mg of protein.

Protein carbonyl (PC) content was measured using the 2,4-dinitrophenylhydrazine (DNPH), according to Levine (Levine et al., 1994). The homogenate pellet was added to 0.5 mL of DNPH solution (10 mmol/L) diluted in hydrochloric acid (7 %), vortexed, and kept at room temperature in the dark, shaking periodically for 30 min. Then, 0.5 mL of ice-cold 10% trichloroacetic acid (TCA) was added to each tube, centrifuged (5,000 g for 10 min at 4 °C), and the supernatant was discarded. The residue was washed three times with 1 mL of ethyl acetate and ethanol (1:1 v/v). Finally, 1 mL of sodium dodecyl sulfate (SDS) 6% was added, the tubes were vortexed, and the supernatant was measured through absorbance at 370 nm. The results were expressed as nmol/ mg of protein based on the molar extinction coefficient of ε370=22 mmol/L × cm.

Total protein was determined according to the Lowry method using bovine serum albumin (BSA) as a standard (Lowry et al., 1951; Waterborg and Matthews, 1994). Total protein concentrations were used to standardize SOD, CAT, GST, MDA, and PC results.

### Statistical analysis

Data distribution was determined by the Shapiro–Wilk test using the program GraphPad Prism 8.0 (San Diego, CA, USA). All data were submitted to unifactorial one-way analysis of variance (ANOVA), followed by Fisher’s Least Significant Difference (Fisher-LSD) post hoc test (α=0.05). Results are expressed as the mean and mean standard error (mean ± SEM). Statistical significance was established at p<0.05.

### Integrated biomarker response (IBR)

To integrate the results from different biomarkers and understand the global response, we calculated the integrated biomarker response (IBR), following the method developed by S. Devin (Devin et al., 2014), using the program CALIBRI (CALculate IBR Interface). The mean (m) and standard deviation (s) of a given biomarker were measured, and the group mean (X) is the mean value for the biomarker for a group. After that, we calculated a standardization for each group to obtain Y:

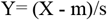

Then, instead of transforming each biphasic biomarker into two variables with positive and negative values relating to biomarker inhibition/activation, we used the square form control Y score:

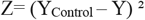

The score results (S) are:

S = Z+|Min|, where S ⩾ 0 and |Min| is the lowest absolute value of Z.

Star plots were then used to display and calculate the integrated biomarker response (IBR), where the IBR is the star plot’s total area. As the results for each organ are a set of biomarkers, a ray coordinate of the star chart represents the score of a given biomarker in a given organ. To avoid the strong dependencies on the biomarker arrangement along the star plot, the program uses a permutation procedure leading to (k−1)!/2 possible values, where k is the number of biomarkers. The final IBR score is a mean of all IBR values corresponding to every possible order of biomarkers along the star plot.

## RESULTS

### Biochemical biomarkers

The treatment with HMs did not affect plasma triglycerides, cholesterol, or glucose levels. However, animals exposed to Cr showed an increase in albumin (p=0.0425), ALT (p=0.0346), and AST (p=0.0187) levels, and animals exposed to Cd increased only the ALT (p=0.0005) when compared with control groups (Fig. 1).

**Figure 1.**
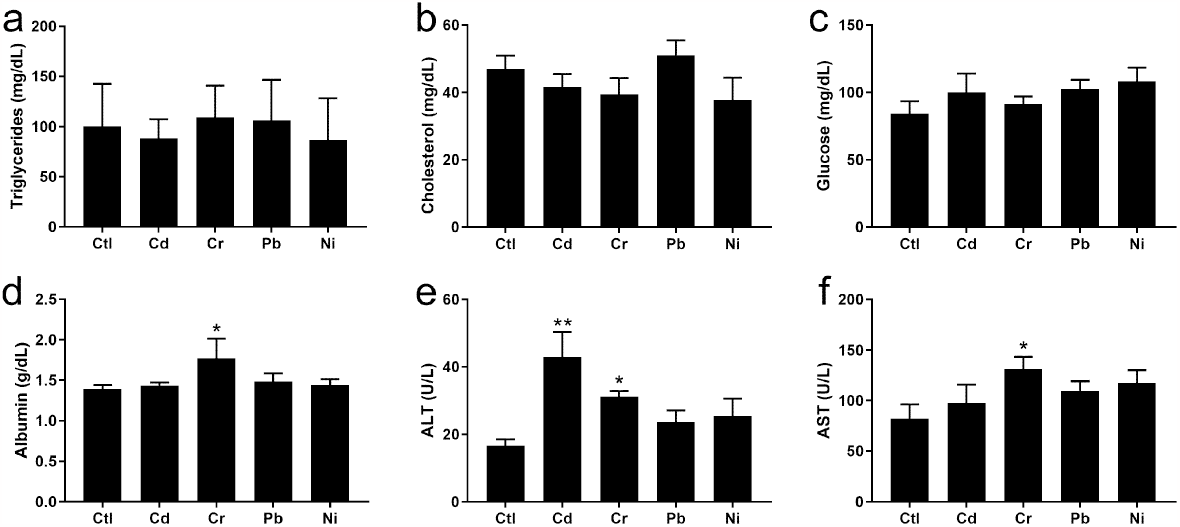
Plasma biochemical markers. (a) Triglycerides, (b) cholesterol, (c) glucose, (d) albumin, (e) ALT, (f) AST. Data are represented as means ± SEM. * p≤ 0.05, ** p≤ 0.01 compared to control group.

### Detoxication biomarkers

In the liver, SOD activity increased only in Ni exposed group (p<0.0001) compared to the CTL group. The catalase activity was not affected by any HMs. However, the GST activity decreased in all exposed groups [Cd (p=0.0010), Cr (p=0.0104), Pb (p=0.0003), and Ni (p=0.0006)] when compared to the CTL group. The FRAP capacity decreased only in Cd treated group (p=0.0165) (Fig. 2).

**Figure 2.**
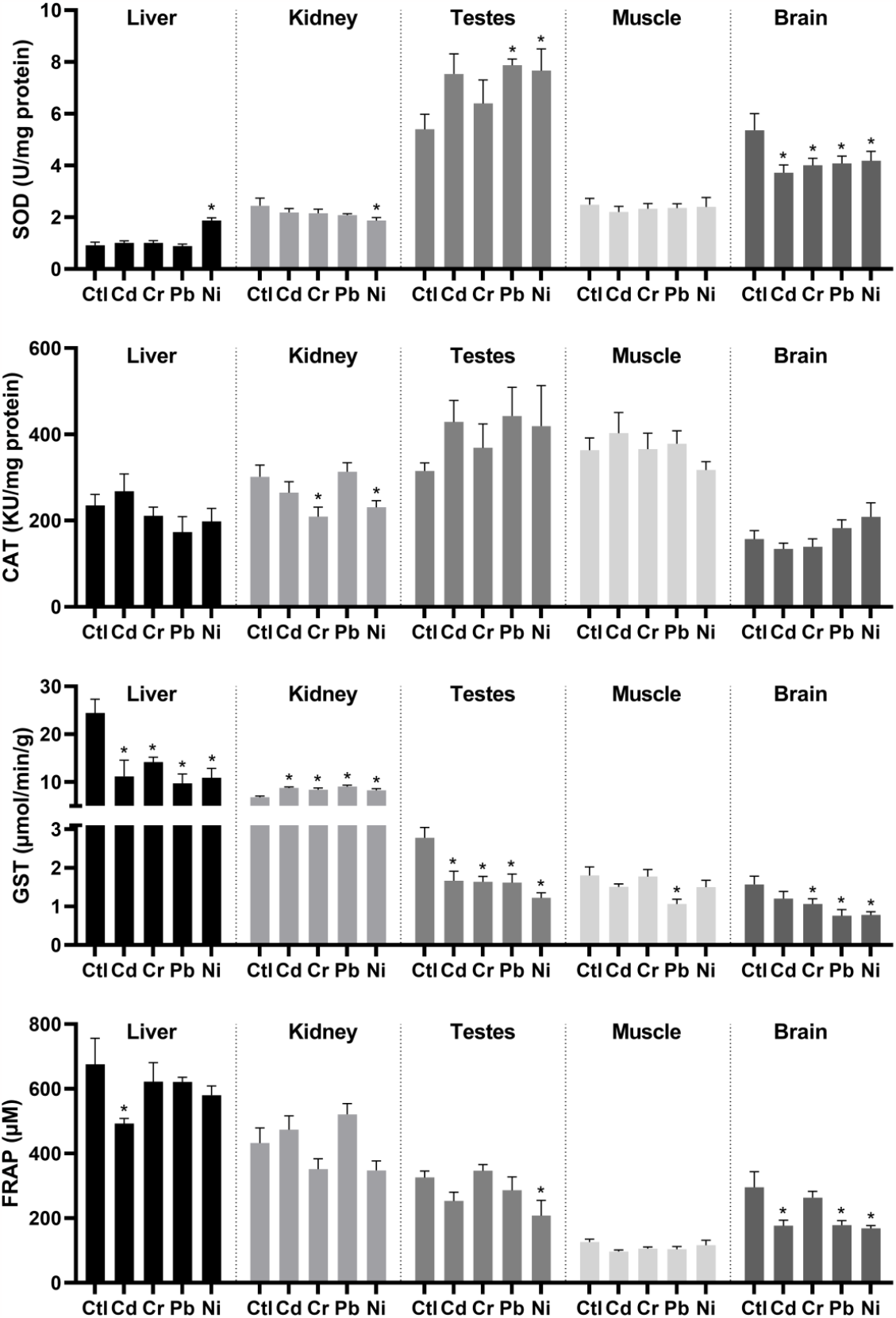
Detoxication biomarkers. The bats were exposed to each heavy metal for 96 hours to 1.5mg/kg (i.p.). The tissues were analyzed for superoxide dismutase (SOD), catalase (CAT), glutathione S-transferase (GST), and ferric-reducing ability of plasma (FRAP). Data are represented as means ± SEM. * p≤ 0.05 compared to the control group.

**Figure 3.**
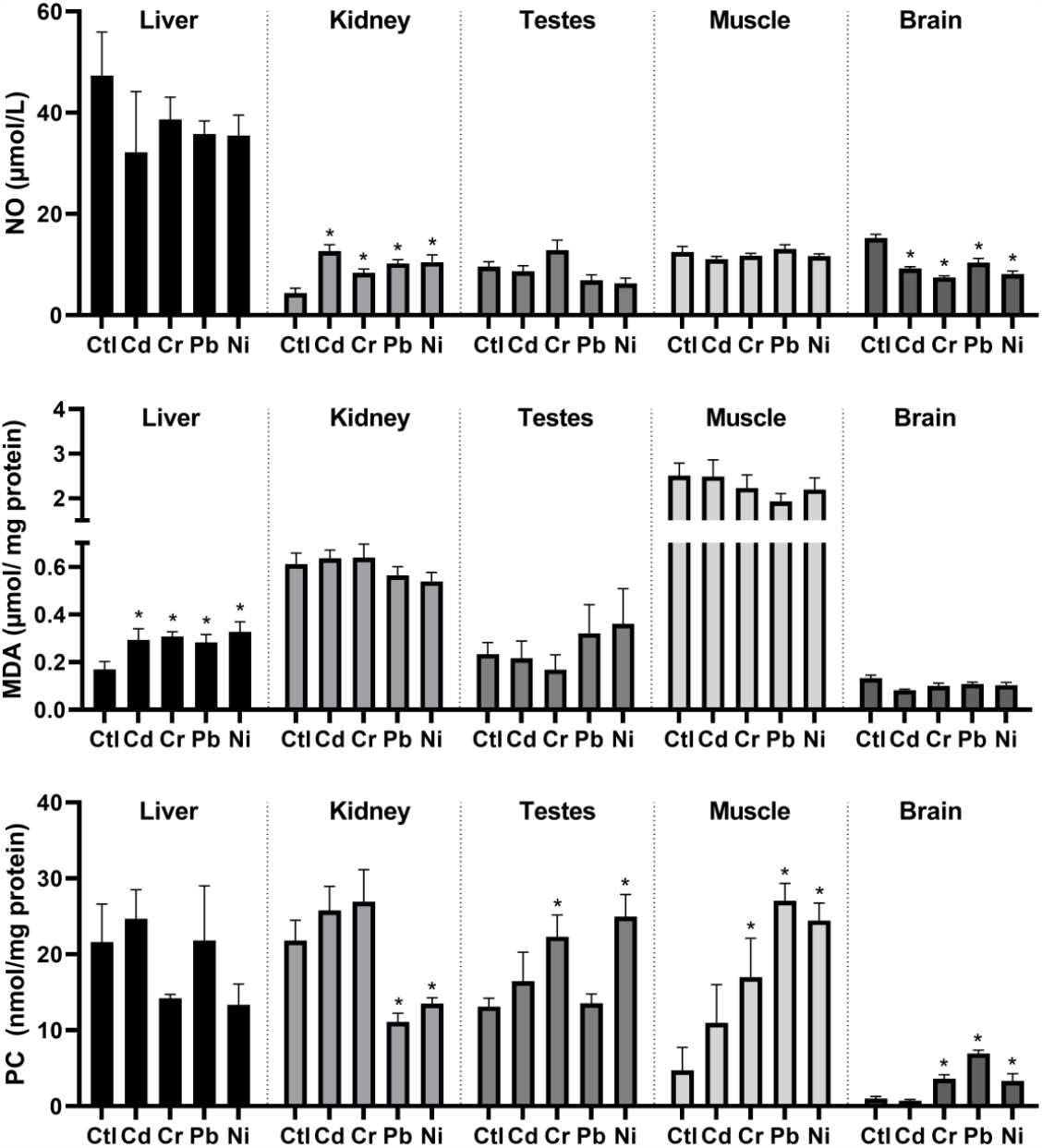
Oxidative damage biomarkers. The bats were exposed to each heavy metal for 96 hours to 1.5mg/kg (i.p.). The tissues were analyzed for nitric oxide (NO), malondialdehyde (MDA), and protein carbonyl (PC). Data are represented as means ± SEM. * p≤ 0.05 compared to the control group.

In the kidney, SOD activity show reduced only in the Ni group (p=0.0214) compared to the CTL group. The catalase activity was decreased in Cr (p=0.0086) and Ni (p=0.0376) groups. However, the GST activity increased in all treated groups [Cd (p=0.0004), Cr (p=0.0018), Pb (p=0.0001), and Ni (p=0.0043)] when compared to the CTL group. The FRAP capacity was unaffected by any HMs (Fig. 2).

In the testes, the SOD activity increased in Pb (p=0.0345) and Ni (p=0.0353) groups. The catalase activity was not affected by any HMs. However, the GST activity decreased in all exposed groups [Cd (p=0.0009), Cr (p=0.0007), Pb (p=0.0009), and Ni (p<0.0001)] when compared to the CTL group. The FRAP capacity is lower only in Ni exposed group (p=0.0209) (Fig. 2).

In the muscle, either SOD, CAT, or FRAP activities were unaffected by any tested HMs. Only the Pb-exposed group showed a significant reduction (p=0.0052) in GST activity (Fig. 2).

In the brain, the SOD activity showed a decrease in all exposed groups [Cd (p=0.0078), Cr (p=0.0243), Pb (p=0.0332), and Ni (p=0.0400)] when compared to the CTL group. However, catalase activity showed no significant increase in any exposed groups. The GST activity showed reduced in Cr (p=0.0327), Pb (p=0.0013), and Ni (p=0.0012) exposed groups. The FRAP capacity decreased in Cd (p=0.0032), Pb (p=0.0035), and Ni (p=0.0018) exposed groups compared to CTL (Fig. 2).

### Damage biomarkers

The exposure to HMs in the liver showed no significant difference in NO production and protein carbonyl levels compared to the CTL group. However, the levels of MDA increased in all exposed groups [Cd (p=0.0235), Cr (p=0.0127), Pb (p=0.0363), and Ni (p=0.0049)] (Fig.3).

In the kidney, the NO showed an increase in all exposed groups [Cd (p<0.0001), Cr (p=0.0313), Pb (p=0.0018), and Ni (p=0.0009)]. Nevertheless, the MDA levels were not affected in any exposed groups compared to the CTL group. Curiously the levels of the protein carbonyl presented a reduction in Pb (p=0.0086) and Ni (p=0.0311) exposed groups (Fig.3).

In the testes, no significant differences are noticed in NO production and MDA levels in all exposed groups compared to the CTL group. The protein carbonyl levels increased in Cr (p=0.0250) and Ni (p=0.0038) exposed groups (Fig.3).

In the muscle, like in the testes, no difference had shown in NO production and MDA levels compared to the CTL group. The protein carbonyl levels increased in Cr (p=0.0250), Pb (p=0.0001), and Ni (p=0.0003) exposed groups (Fig.3).

In the brain, the production of NO had impaired in all exposed groups [Cd (p<0.0001), Cr (p<0.0001), Pb (p<0.0001), and Ni (p<0.0001)]. However, the MDA levels were not affected in any HM exposed groups compared to the CTL group. The protein carbonyl levels increased in Cr (p=0.0049), Pb (p<0.0001), and Ni (p=0.0091) exposed groups (Fig.3).

### Integrated biomarker response (IBR)

The IBR star plots of the liver, kidney, testes, muscle, and brain were shown (Fig. 4).

**Figure 4.**
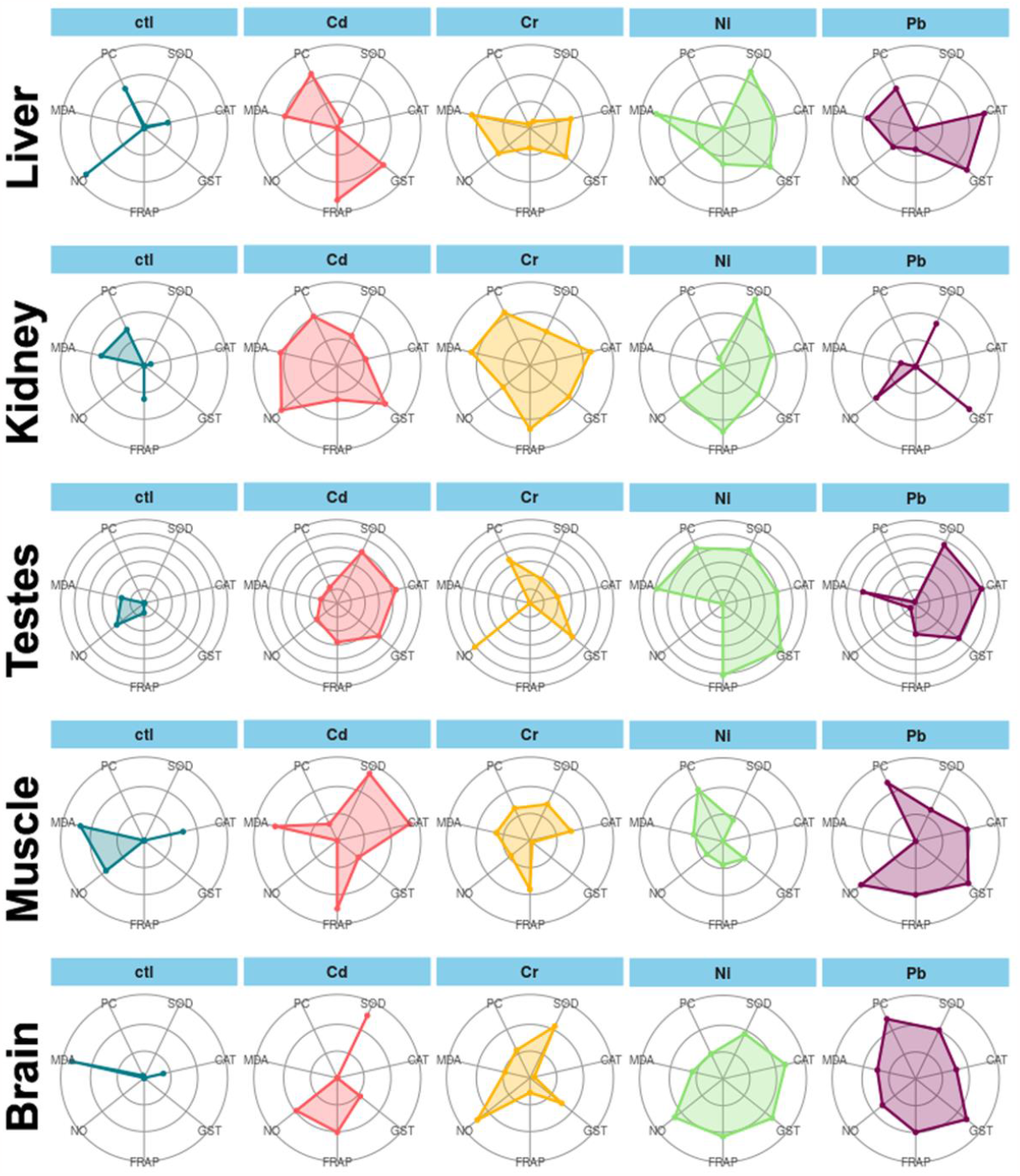
Star plot of tissues exposed to heavy metals. The total area of star plots represents the value of the IBR index.

All groups exposed to HMs in the liver, testes, and brain had a rise in global damage. In the kidney, the groups exposed to Cd, Cr, and Ni had global damage increased. In muscle, the global damage increased in groups exposed to Cd, Cr, and Pb when compared to the CTL group (Fig. 5).

**Figure 5.**
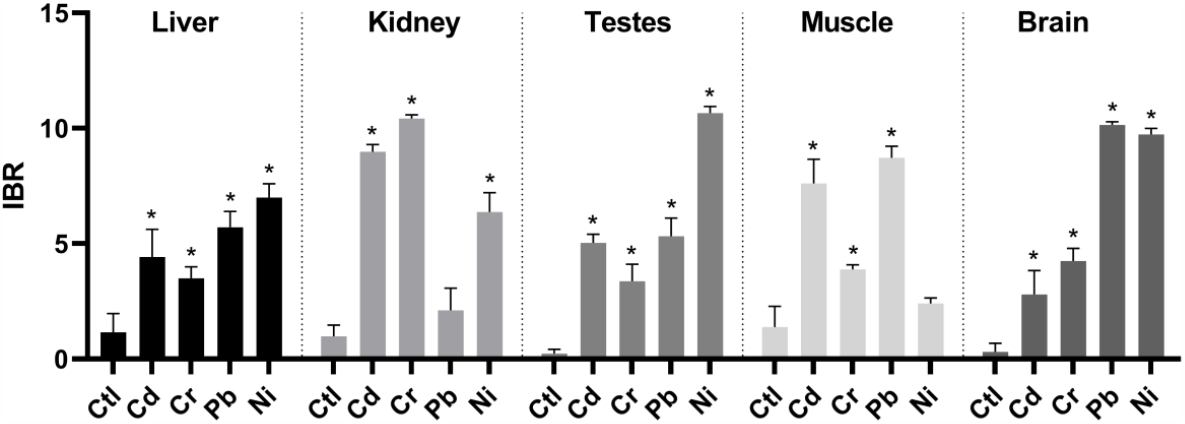
The integrated biomarker response (IBR) index. Global analysis of either detoxication and damage biomarkers. Data are represented as means ± SEM. * p≤ 0.05 compared to the control group.

## DISCUSSION

Some metals are essential to physiological processes in animals. Still, excessive environmental concentrations through anthropic activities may potentially impair reproduction and other functions that could lead to decreases in wildlife populations in the long term (Karri et al., 2016). This is the first study evaluating the isolated effect of HMs on the oxidative status of wild bats using the IBR index. The concentrations analyzed in this study are lower than the concentrations found in insectivorous bats living in coal mining areas (José Zocche et al., 2010) and lower than those found in tropical basin sediments in Brazil, so they are considered low concentrations (Quadra et al., 2019). Variable concentrations of Cd, Cr, Pb, and Ni are often found, depending on the proximity of industrial activities and the soil’s natural composition. For instance, concentrations (mg/kg) up to 11.7 for Cd, 468 for Cr, 510 for Pb, and higher than 583 for Ni in the soil close to gold mining exploration have already been reported (Fashola et al., 2016).

### Biochemical markers

Were observed that the used concentration of 1.5 mg/kg of each HM had little or no influence on triglycerides, cholesterol, glucose, albumin, ALT, and AST. The metals Pb and Ni unexpectedly did not influence any biochemical markers. Cd induces an increase only in ALT levels. Cr was the metal that caused little but significant alterations in biochemical biomarkers, increasing albumin, ALT, and AST. Together with all results, the biochemical markers showed to be not so sensitive to predict the damage caused by HMs.

### Liver alterations

The global analysis of IBR in the liver demonstrates that exposure to all metals harms the organ. However, the liver looks more sensitive to Ni and Pb than Cd and Cr. The antioxidant enzymes were differently influenced by Cd, Cr, Ni, and Pb. SOD activity was increased only by Ni, and FRAP was decreased only in Cd exposed group. However, all metals halved the GST activity.

The physiological role of Ni in vertebrates is still not well understood. Ni is an essential element in plants, bacteria, and animals. Its deficiency affects the metabolic processes, increasing total lipids in the liver and decreasing phospholipids (Mishra and Singh Sangwan, 2019). Here, Ni exposure increased SOD activity, which may be associated with an overproduction of hydrogen peroxide (H_2_O_2_) (Ighodaro and Akinloye, 2018). Excessive H_2_O_2_ formation would further activate CAT and Glutathione peroxidase (GPx). Our study showed no increase in CAT activity following HMs exposure. GST activity decreased, indicating this enzyme could no longer protect the cells from lipid peroxidation (Recknagel et al., 2020), corroborating the increased MDA observed in all groups. The decreased GST activity might be explained by this metal’s ability to bind to thiol (-SH) groups on proteins and make them inactive (Balali-Mood et al., 2021). We suggest that this decreased activity of GST also decreases GSH activity since GST catalyses the conjugation of GSH to toxic electrophiles (Zhang et al., 2021). Corroborating this, a decrease in GSH activity and sulfhydryl content was observed in lungs from *Artibeus lituratus* collected in a mining area (Pedroso-Fidelis et al., 2020). Environmental studies showed that insectivorous bats living in coal mining areas could bioaccumulate higher metal concentrations in the liver than those analysed in our research. These bats showed DNA damage in blood cells (José Zocche et al., 2010) which may be correlated with changes in their redox status.

### Kidney alterations

The global analysis of the kidney showed us an unexpected result the amount of Pb used in the study was not harmful to the kidney. However, the exposure to other metals (Cr, Cd, and Ni) was toxic. The antioxidant enzymes were influenced by Cd, Cr, Ni, and Pb, in different ways. All these slight alterations appear no has biological importance. The most critical result observed in the kidney has increased NO production induced by metal exposure. Low levels of metals in chronic exposure lead to increased blood pressure, eNOS/iNOS in the kidney, and reduced urinary NO excretion (Lee, 2008). NO has several kidney functions regulating renal hemodynamics and tubuloglomerular feedback (Mount and Power, 2006). NO produced by the macula densa in the kidneys inhibits tubular sodium reuptake, resulting in increased urinary excretion and water and solutes lee, contributing to the excretion of toxic compounds. NO reacts with superoxide radical (O^2-^) and produces peroxynitrite (ONOOH^-^), which can produce hydroxyl radicals (HO^•^) in the presence of hydrogen ions (Oliveira et al., 2017).

Therefore, an increase in NO without a sufficient increase in antioxidant enzymes can lead to cellular oxidative stress.

### Testes alterations

Regarding the IBR index, all HMs presented a global increase in damage levels, highlighting the Ni that doubled compared to the other metals. The antioxidant defence of the testes presents an activity profile like the liver, where GST was negatively affected by all HMs exposure. GSTs are a family of antioxidant isoenzymes that participate in the cellular detoxification of several xenobiotics. The inhibitory effects of metals in GST activity may be harmful to the cells (Guven and Soydan, 2022). Several studies showed decreased GST activities following HMs exposure in testes of mammals, such as decreased GST after 20mg/kg of Pb (i.p.) for 5 days in rats (Moniem et al., 2010), after 0.025mg/kg of Pb and Cd (ipi) for 15 days in rats (Pandya et al., 2012) and after a single dose of 3.58 mg/kg of Cd and 59.5 mg/kg of Ni (i.p.) in rats (İşcan et al., 2002). Here, we found that exposure to Ni and Cr induced protein carbonyl in male gonads. Protein carbonylation is an irreversible post-translational modification caused by excessive ROS formation, considered the main hallmark of oxidative stress disorders (Fedorova et al., 2014). Overall, the testes look more sensitive to Ni than other metals.

### Muscle alterations

The global analysis of IBR in the muscles demonstrated that Ni is not harmful to this tissue. However, the other metals (Pb, Cd, and Cr) were toxic to the muscles, which look more sensitive to Pb and Cd than to Cr exposure. Cd, Cr, Ni, and Pb poorly influenced the antioxidant enzymes. Only GST shows a slight decrease in Pb exposed group.

Pb, Ni, and Cr induced more oxidative damage in muscles than Cd by protein oxidation. Bats’ breast muscles are intensively used during flight. The high O_2_ offer needed to address high metabolic activities (Costantini et al., 2019) usually involves high ROS production, demanding an efficient antioxidant capacity from these tissues. We found that Pb exposure decreases the enzymatic activity of the antioxidant enzyme GST, which may culminate in protein carbonylation. This oxidative damage can be detrimental to bat flight.

### Brain alterations

Although the blood-brain barrier (BBB) protects brain tissue, toxic metals with similar size and chemical properties of essential metals can cross the barrier (Hill et al., 2018). The global analysis of the IBR index showed that exposure to all metals was harmful to the brain.

However, the brain looks more sensitive to Pb and Ni than Cr and Cd. The brain was also sensitive regarding the antioxidant ability in metals-exposed bats. Since SOD was impaired in all exposure groups, GST was decreased in Cr, Pb, and Ni groups, and FRAP was reduced following Cd, Pb, and Ni exposures. Corroborating our results, bats captured near a water treatment site had decreased total antioxidant capacity and high levels of arsenic and Pb in the brain (Pb concentrations were close to the value analysed in our experiment (1.169 mg/kg)) (Hill et al., 2018).

In bats’ brains, NO levels decreased in all exposed groups. The inhibitory effect of HMs on nitric oxide synthase (NOS) activity in mammals has been documented (Mittal et al., 1995). In this study, exposure to Cr, Pb, and Ni induced cell damage, especially protein oxidation. Rats exposed to higher doses of Pb (50mg/kg for 5 days) also showed damage in proteins (Reckziegel et al., 2011). Pb is known to interfere with Ca^2+^ signalling pathways in the hippocampus, a brain region associated with memory and learning (Karri et al., 2016).

Therefore, Ca^+2^-regulated neuronal processes, such as NO synthesis, might be affected, which could impair cognitive abilities related to foraging, echolocation, and flight capacities (Hill et al., 2018; Karri et al., 2016). Behavioural changes associated with alterations in the circadian cycle have been observed in insectivorous bats captured in areas exposed to HMs (Lovett and McBee, 2015).

Although they were all captured from the same protected area, they might have been exposed to other pollutants before the experiment. Experiments carried out with wild bats have limitations regarding the origin of these animals. Captured animals do not have the same controlled background as laboratory animals, born and raised under similar conditions.

However, we used the control group to compare animals in the same condition, sex, and location. Other experiments run by our team turned out to be successful in bringing together scientific subsidies to understand better the effects of pollutants on bats (Brinati et al., 2016; Freitas et al., 2021; Oliveira et al., 2017; Oliveira et al., 2021). We kept animals under an acclimating period before the experiment to reduce bias. Also, all bats were captured in the same location and during the same season in the same year. Furthermore, we propose an order of concern for metals pollution based on the sum of IBR indexes of all analysed organs in bats: Ni > Pb > Cd > Cr.

Taken together, our results draw attention to possible oxidative and tissue damage from HMs exposure in fruit bats, even under a short time. Our results indicate that bats may be susceptible to HMs’ effects. In addition, we propose the following order of metal toxicity based on the IBR indexes: Ni> Pb> Cd> Cr. These findings demonstrate that HMs, constantly released into the environment, may affect bats’ reproductive and cognitive capacity and ecological contribution. Future studies with different concentrations and exposure times will help understand other toxicological effects better. Assessing and monitoring bat populations in highly contaminated areas are critical to know better the damage caused by environmental pollution to key ecological species.

## Acknowledgements

The Brazilian agencies supported this work: Fundação do Amparo à Pesquisa do Estado de Minas Gerais (FAPEMIG) and Conselho Nacional de Desenvolvimento Científico e Tecnológico (CNPq). Destro received a PhD fellowship from CAPES (Coordenação de Aperfeiçoamento de Pessoal de Nível Superior). LL Oliveira and RV Gonçalves are CNPq Research Productivity fellows.

## Authorship contributions

**Ana L. F. Destro**: Writing − original draft, Investigation, Sampling, Data analysis. **Thaís S. Alves**: Sampling, Data analysis. **Fernanda R. Dias**: Sampling, Data analysis. **Reggiani V. Gonçalves**: Resources, Software, Investigation. **Jerusa M. Oliveira**: Investigation, Writing - original draft. **Leandro L. Oliveira**: Resources, Software, Writing − review & editing. **Mariella B. Freitas**: Writing, Conceptualization, Review, Editing, Supervision, Funding acquisition. All authors read and approved the manuscript.

## Ethical Approval

All animal captures and procedures performed in this study were approved by the Brazilian Government (SISBIO 75064-1) and by the Animal Ethics Committee (CEUA-UFV 26/2020).

## Data availability

The datasets used and/or analysed during the current study are available from the corresponding author upon reasonable request.

